# Sex differences in latent inhibition and their implications for psychotic disorders

**DOI:** 10.1101/2025.09.24.677991

**Authors:** Eleanor R. Mawson, Jeremy Hall, Kerrie L. Thomas

## Abstract

Latent inhibition (LI) refers to the reduced ability to learn or express learning about a stimulus following prior exposure to it without consequence. It has been extensively studied to investigate associative learning and memory as well as the aberrant attribution of salience to irrelevant stimuli in both humans and animals, a process which is thought to underpin hallucinations and delusions in psychosis. Despite sex differences observed in numerous aspects of psychosis, few studies have attempted to characterise sex differences in LI. This systematic review outlines the current evidence of sex differences in LI in both human and rodent studies, showing predominantly inconclusive and conflicting evidence for sex differences. However, the majority of cases in which sex differences were found were in studies that assessed the impact of additional factors on LI such as drug administration, juvenile handling, and oestrous phase, and demonstrated interactions between sex and these additional factors. Hence, there is stronger evidence for sex differences in vulnerability to perturbations in LI. Furthermore, we present data demonstrating sex differences in latent inhibition of contextual fear conditioning and associated gene expression in the hippocampus in rodents, suggesting sex differences in hippocampal processing underpinning latent inhibition at both the behavioural and neurobiological levels. Although there have been relatively few studies demonstrating sex differences in LI identified by the systematic review, the data suggest that in hippocampally dependent forms of LI, sex differences may be more apparent, which may be of relevance in relation to sex differences in the presentation of psychosis.

## 1. Introduction

Associative learning is one means through which humans and other animals utilise knowledge of prior experiences that have co-occurred with a stimulus or environment to predict future outcomes and direct behaviour. Latent inhibition (LI) refers to the reduced behavioural expression of associative learning and memory produced by the pairing of a conditioned stimulus (CS), or context (i.e., the environment in which learning takes place), with an innately rewarding or aversive unconditioned stimulus (US) when the subject has been previously exposed to the CS, or context, alone without any reinforcement[1]. LI is observed in both humans and animals, and the majority of both rodent and human conditioning experiments have paired a discrete CS with a US, a paradigm known as cued conditioning[2]. Context plays an important role in the expression of LI in cued conditioning, and LI is typically observed only when testing occurs in the same context in which pre-exposure and conditioning took place[3]. Conditioning in the absence of a discrete cue, typified by contextual fear conditioning (CFC) paradigms, leads to a direct association between the context and the US, sometimes termed as foreground contextual conditioning[4]. Latent inhibition experiments employing CFC protocols involve pre-exposing the individual to the context prior to fear conditioning[2]. In humans, deficits in LI have been implicated in psychosis[5,6]. Sex differences are seen in the clinical presentation of psychosis, such as age of onset, symptom presentation and response to treatment[7]. Despite the extensive body of research on LI, most studies have not included sex as a factor in their analysis and behavioural neuroscience studies have been historically biased towards the sole inclusion of male animals. Consequently, important sex differences in cognitive mechanisms underpinning psychosis that have the potential to explain sex differences in clinical presentation may have been overlooked. Furthermore, when developing novel precision treatments that target underlying biology, if this biology differs between sexes, treatment efficacy may be unequitable between sexes.

This review aims to outline our current understanding of the sex differences in LI by conducting a systematic review of the literature in both rodents and humans across both cued and contextual associative learning paradigms. Furthermore, novel data are presented relating to sex differences in the LI of contextual fear conditioning in rats. Additionally, the relationship between LI and psychosis will be explored, and how characterising sex differences in LI can be used to improve our understanding of sexually dimorphic psychopathologies such as schizophrenia and bipolar disorder will be discussed.

## 2. Mechanisms underpinning Latent Inhibition

Several theories have attempted to explain the phenomenon of LI whereby nonreinforced preexposure of a CS results in impaired acquisition or expression of subsequent CS-US associations. Attentional theories such as the Conditioned Attention Theory[8,9] posit that LI functions as an adaptive mechanism to direct limited attentional resources to stimuli that best predict relevant outcomes, thereby conserving cognitive resources for novel or predictive stimuli. Conversely, associative or comparator theories assert that LI occurs as a result of competing CS-no US or CS-context associations formed during pre-exposure and CS-US acquired during conditioning[6,10–13].

Dopamine plays a central role in attentional control[14], context processing[15], and associative learning processes[16] important for LI. Administration of both typical and atypical D_2_ receptor antagonist antipsychotics induce enhancement of LI[17–21]. Conversely, administration of amphetamine has been shown to inhibit LI effects in rodents[22]. Attentional theories would predict that the effects of antipsychotics and amphetamines would be exerted during the pre-exposure period. However, such effects have been shown to be specific to administration during the conditioning phase[23,24]. Thus, this evidence would suggest that dopaminergic signalling does not wholly underpin the learning about irrelevant (unreinforced) stimuli but at least partially exerts its effects on the utilisation of this information during conditioning.

Converging evidence from lesion, pharmacological, and optogenetic studies has implicated a distributed neural circuit in LI, including the ventral tegmental area (VTA), nucleus accumbens (NAc), medial pre-frontal cortex (mPFC), and hippocampus. A wide range of evidence supports the involvement of dopaminergic signalling in this network. Amphetamine administration as well as more recent optogenetic experiments have revealed that dopaminergic input from the VTA to the NAc plays a key role in LI, with dopamine activity in the NAc inversely related to LI expression[25–27]. Furthermore, amphetamine administration potentiates dopaminergic signalling from the mesolimbic pathway to the NAc and wider ventral striatum[28], and, as is previously mentioned, this has been shown to attenuate the expression of LI in rodents[22].

The hippocampus is integral to the formation and retrieval of conjunctive representations of environmental contexts[29,30]. Accordingly, the hippocampus is critically involved in contextual learning, including the formation and retrieval of contextual fear memories and their associated stimulus–outcome relationships[31]. Context is important for LI because LI is diminished when pre-exposure and subsequent conditioning or testing occur in different contexts[12,32], and the hippocampus has been shown to be crucial to this context-dependent specificity[33]. During pre-exposure, the animal is therefore thought to form a representation of the surrounding context, which can subsequently modulate the processing and associative significance of the pre-exposed stimulus. The hippocampus also forms part of a functional circuit with the VTA and dopaminergic systems that has been proposed to facilitate the encoding of novel information into long-term memory[34]. The entorhinal cortex (ENT) interfaces with the hippocampus and regulates dopaminergic transmission in the NAc, providing a potential pathway through which contextual and novelty-related information may influence mesolimbic dopamine signalling[35].

Alongside the hippocampus, the IL and PL regions of the mPFC appear to play distinct and potentially competing roles in regulating the context specificity of LI. The importance of the mPFC to LI has been shown through the use of functional activation and inactivation experiments. Nonlocal lesions or amphetamine infusions into the mPFC have no effect on LI[36,37]. However, inactivation specifically of the IL disrupts LI, indicating a general function for the IL in the formation and retrieval of CS-no US and context-no US associations[38,39]. Conversely, catecholaminergic depletion of the PL has been shown to enhance LI, indicating a distinct role for the PL in regulating its expression[40]. Consistent with this regional specificity, excitotoxic lesions of the ventral/IL, but not dorsal/PL mPFC have been shown to enhance LI[41]. Recent studies suggest a more nuanced role for mPFC regions, with the PL and IL modulating the retrieval of initial CS-no US and subsequent CS-US associations, respectively, such that manipulations of the PL or IL can inhibit or enhance LI[41,42]. Alterations in dopaminergic signalling in the NAc and PFC, alongside altered transmission of glutamate and GABA, have been implicated in the neurobiology underpinning psychosis[43]. Hence, studying the role of dopamine in cognitive processes thought to relate to psychosis, such as LI, may provide insight into the mechanisms underpinning aberrant salience attribution in psychosis.

## 3. Latent Inhibition and Psychosis

Psychosis is a frequently debilitating clinical phenomenon characterised by hallucinations and/or delusions and is a hallmark feature of disorders such as schizophrenia (and schizophrenia spectrum disorders) and is commonly seen in other conditions such as bipolar disorder[44]. Deficits in LI have been observed in schizophrenia[5,6], and consequently LI has been studied in the context of psychosis in numerous studies to model the underlying cognitive mechanisms[45–47]. It has been hypothesised that LI disruptions occur in psychosis owing to altered dopaminergic signalling. Increased striatal dopamine synthesis capacity and synaptic dopamine content have been observed in psychosis, with the striatum receiving input from a wider network of the thalamus, hippocampus and prefrontal cortex[48].

The aberrant salience hypothesis of schizophrenia argues that directed attentional processes are dysfunctional in schizophrenia and alterations in dopaminergic signalling lead to misdirection of attention to irrelevant stimuli, resulting in hallucinations and delusions[47,49]. Prediction error and computational Bayesian theories explaining the altered dopaminergic signalling observed in psychosis assert that this dysfunctional dopaminergic signalling leads to prediction errors encoding irrelevant stimuli as salient, i.e., aberrant salience[50,51]. The dopamine-driven prediction of non-salient events as surprising results in heightened value attributed to such stimuli, leading to the positive symptoms of psychosis such as hallucinations and delusions. Conversely, the reduced phasic dopamine response for relevant stimuli may underpin the development of negative symptoms seen in schizophrenia, such as anhedonia and blunted affect[52]. Thus, LI deficits in psychosis may arise from altered dopaminergic signalling resulting in previously pre-exposed irrelevant stimuli having inappropriately high associability during learning due to disrupted prediction error processing. Other theories, such as the switching model, suggest that disrupted dopaminergic signalling results in heightened attentional switching between stimulus–no outcome associations learned during pre-exposure and stimulus outcome associations learned during conditioning to control behavioural response, supported by evidence that heightened dopamine increases cognitive switching and antipsychotics have been shown to reduce cognitive switching[53]. Thus, LI occurs as a result of heightened cognitive switching between the CS-no event association acquired during pre-exposure and the CS-reinforcement association acquired during conditioning[27].

Studies of LI in the context of psychosis show LI to be attenuated in people with schizophrenia and those scoring highly on measures of schizotypy, albeit with some variability between studies. However, most do not include sex as a biological variable in analyses, even when studies are conducted with both male and female participants[5]. This is despite the large body of literature that has revealed sex differences in psychosis in schizophrenia and bipolar disorder. Schizophrenia has a higher prevalence[54] and earlier age of onset[55] in male compared to female individuals. Differences have also been observed between sexes in relation to pattern of symptoms, number of psychotic episodes, and comorbid psychopathologies[55,56]. In healthy individuals, subclinical psychotic experiences appear to be more prevalent in male than female individuals during adolescence, but by early adulthood this sex difference disappears[57]. However, middle age represents a risk period for psychosis development in women, and women make up the majority of diagnoses at or post middle age, a finding which is thought to be attributable to hormonal changes occurring during the menopause[58]. Despite the increasing recognition of sex as an important biological variable in neuroscience, many studies investigating the biological and cognitive mechanisms relevant to psychosis have historically failed to examine sex differences[59]. Similarly, a recent review of LI in schizophrenia and schizotypy found that, sex differences were not systematically examined in the majority of studies[5]. Hence, this systematic review evaluated sex differences in LI studies where both sexes were included in both human and rodent (rat and mouse) experiments.

## 4. Systematic Review of Sex Differences in Latent Inhibition

### 4.1. Methods

#### 4.1.1. Search strategy

“Latent inhibition” was entered as a search term into SCOPUS, Web of Science, PUBMED and Science Direct databases on 15^th^ September 2024, with all results retained for screening. No restrictions were placed on the year of publication. Although LI in humans has been studied largely in the context of psychosis, this review aimed to assess any LI study in humans regardless of clinical status of cohort. Subsequently, any studies in cohorts of individuals with psychosis (e.g, schizophrenia, bipolar disorder or schizoaffective disorder), clinical cohorts of individuals with other psychiatric conditions, and any non-clinical cohorts were assessed, including studies investigating the impact of schizotypy on LI in non-clinical cohorts. Schizotypy refers to a multi-dimensional personality construct rather than a clinical diagnosis, characterised by cognitive-perceptual (e.g., odd beliefs, hallucinations), negative (e.g. anhedonia, alogia) and disorganised (e.g., thought disturbances, disorganised actions) experiences[60]. High schizotypal individuals have been shown to be at increased risk for developing psychosis, and studying schizotypy in non-clinical cohorts allows for a wider exploration of the aetiology of the disorder[61].

It is important to note that although sex and gender are distinct constructs, the terminology of gender and sex have largely been used without distinction within the studies reviewed. In this review, the term ‘sex’, with the binary of ‘male’ and ‘female’ will be used to refer to human participants, as this binary categorisation has been used within the literature, irrespective of whether the term ‘sex’ or ‘gender’ has been used. It is acknowledged that this categorisation does not correspond to the full spectrum of gender, hence the term gender is not used here. Those identifying as a gender that does not correspond to their assigned sex at birth, or are intersex, may have been included in studies reviewed here based on their gender identity rather than their assigned sex at birth, however this is expected to affect only a very small number of participants.

#### 4.1.2. Screening and selection of eligible studies

Results were filtered to remove entries that were not journal articles, and any articles not in English, as well as any duplicated studies. All studies were then screened to identify those that included both male and female participants/animals. Studies were then assessed for eligibility based on the following criteria: a) to only include those that compared a CS or context pre-exposure condition with a non-pre-exposure condition (either between- or within-subject); b) LI independently assessed in males and females, or LI is statistically compared between sexes; c) LI was not assessed using a conditioned taste aversion (CTA) paradigm, as in this unique method the US does not arise until after a considerable delay from presentation of the CS, and the neurobiological underpinnings of CTA are thought to be distinct from other forms of classical conditioning[62].

#### 4.1.3. Data extraction

If a comparison between pre-exposure conditions was used in the study (i.e., pre-exposed vs non-pre-exposed conditions), the interaction between sex and PE condition was assessed. Alternatively, in studies using a metric for LI, such as a ratio or score as the dependent variable, the main effect of sex on this variable was assessed. The interaction between these effects and any further variables were also assessed and reported below. No metanalysis of data was conducted as this review is qualitative in nature.

### 4.2. Results

#### 4.2.1. Systematic review

8908 records were returned after the initial search of the four data bases. After removing any items that were not in English, were not journal articles, or were duplicate items, 4070 records remained. These were screened to only retain records that were studies in either humans, rats or mice that included both male and female participants/animals, resulting in 1057 studies. The following inclusion criteria were used to retrieve the final set of studies: a) a direct comparison between a CS or context pre-exposure and a non-pre-exposure condition (either within- or between-subject); b) LI independently assessed in males and females, or LI is statistically compared between sexes; c) LI was not assessed using a CTA paradigm. 27 studies using human participants and 54 studies using rats or mice remained. The results of the review process are outlined in Fig.1.

**Fig. 1.**
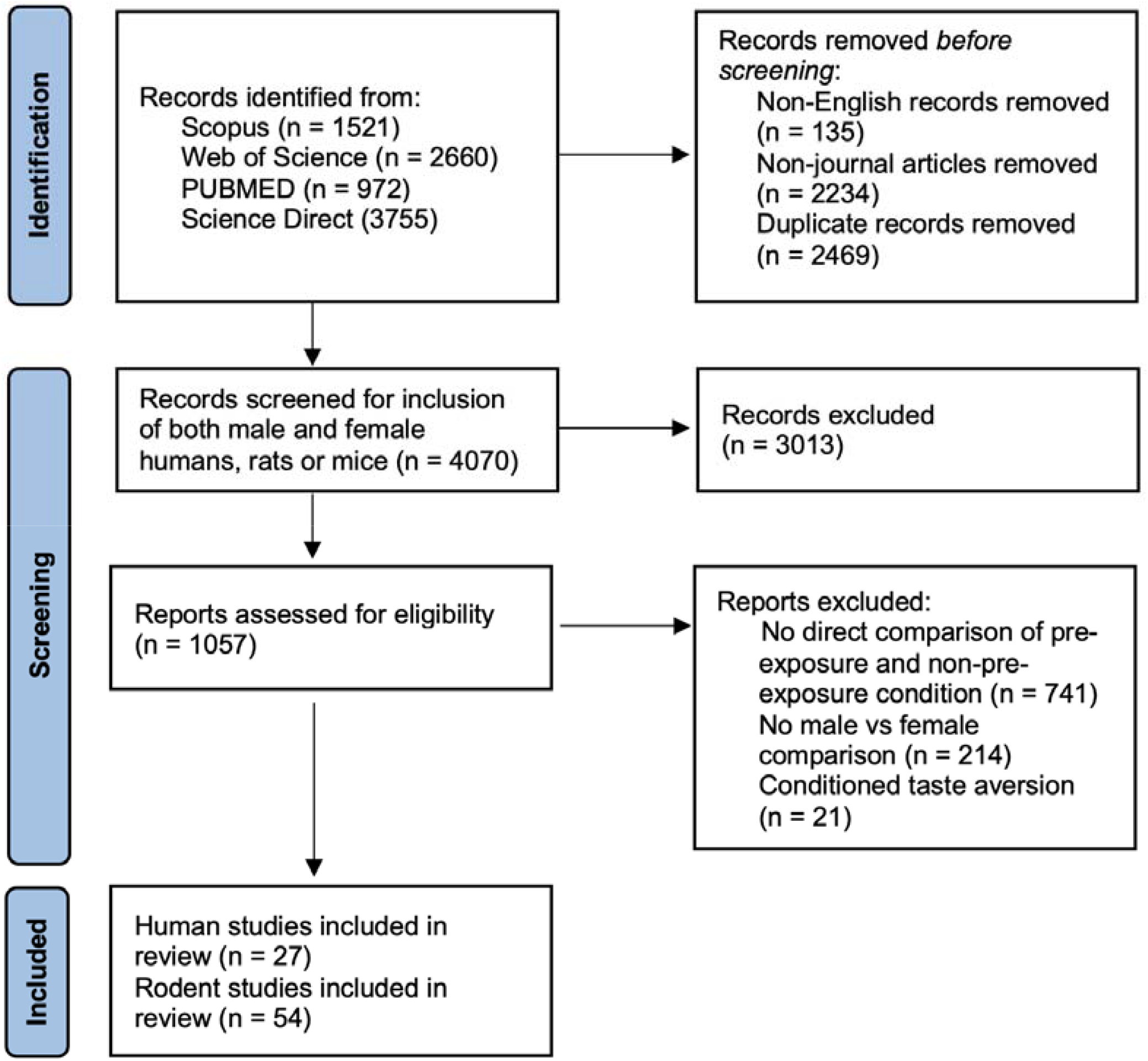
Systematic review process.

#### 4.2.2. Review of human studies

31 experiments across 27 separate studies were found (Table 1) and used either visual or auditory LI tasks with either response reaction time, number of correct responses, or trials to learning criteria as the dependent variable. In all designs, participants learn to use a cue to predict a target. In visual LI tasks, either shapes or letters were used as the CS, and the US may be an aversive noise or token reinforcer such as accumulated points. In the pre-exposure phase, the cue is repeatedly presented on screen with no target. This is followed by a conditioning phase, in which the cue is consistently paired with the target (i.e., the cue reliably precedes and predicts target onset), establishing the cue-target association. In the test phase, participants are instructed to press a button when the target appears, and the reaction time to identify the target is recorded as the dependent variable. In trials to learning criteria designs, cues are similarly pre-exposed depending on the condition, but the conditioning and test phases are typically merged: participants indicate via button-press when they expect the target to appear, and learning is inferred directly from these trial-by-trial anticipatory responses. The number of trials required to achieve a set number of correct responses is used as the dependent variable. In within-subject designs, one cue is presented during the pre-exposure phase (pre-exposed cue), and another cue is only presented during the test phase (non-pre-exposed cue).

**Table 1.** Sex differences in latent inhibition in humans. Studies highlighted in orange show an overall effect of sex on LI (interaction between sex and pre-exposure, or main effect of sex on LI). Studies highlighted in blue show an effect of sex on the relationship between LI and an additional variable (three-way interaction between sex, pre-exposure, and additional variable, or two-way interaction between sex and an additional variable on LI). Conclusions relating to sex differences in LI in each study are highlighted in bold. PE: pre-exposure.

| Year | Authors | Participants | Pre-exposure Effect | Other Variables Measured | Sex Effects on LI |
| --- | --- | --- | --- | --- | --- |
| **Studies in non-clinical adult cohorts** |  |  |  |  |  |
| 2000 | Peterson and Carson[63] | Adults (university students) | Increased trials to criterion in PE compared with no-PE group. | IQ: Positively correlated with PE but not no-PE response.<br>Age: no main effect.<br>Openness to experience: negative correlation with LI score. | No main effect or interaction.<br>**No sex difference in LI.** |
| 2001 | Lubow, Kaplan and De la Casa[64] | Adults (university students) | Reduced reaction time in PE compared with no-PE group |  | No main effect or interaction<br>**No sex difference in LI.** |
| 2001 | Serra, Jones, Toone and Gray[65] | Adults. | Increased trials to criterion in PE compared with no-PE group. | Age/IQ: no main effects or interactions. | No main effect or interaction<br>**No sex difference in LI.** |
| 2002 | Peterson, Smith and Carson[66] | Adults (university students) | Not specified – participants split into high and low LI scorers for subsequent analysis. | Personality variables: Openness to experience, extraversion, plasticity and creative personality associated with lower LI. | No main effect or interactions.<br>**No sex difference in LI.** |
| 2004 | Burch, Hemsley and Joseph[67] | Healthy adults | Visual task:<br>Increased trials to criterion on max-PE and med-PE compared with no-PE group. | Task order/IQ: no main effects.<br>Unusual experiences: LI disrupted in high unusual experiences group. | No sex difference in LI, or interaction with unusual experiences.<br>**No sex difference in LI.** |
|  |  |  | Auditory task:<br>Increased trials to criterion in max-PE, med-PE and min-PE compared with no-PE groups. | Task order/IQ: no main effects.<br>Unusual experiences: main effect not reported. | No sex difference in LI, or interaction with unusual experiences.<br>**No sex difference in LI.** |
| **Studies assessing schizotypy in non-clinical adult cohorts** |  |  |  |  |  |
| 1997 | Williams, Wellman, Geaney, Feldon, Cowen and Rawlins[68] | Healthy adults | Greater trials to criteria in PE compared with no-PE group. | Haloperidol/schizotypy/smoking status: no main effects. | No interactions.<br>**No sex difference in LI or interaction with haloperidol.** |
| 1999 | De la Casa, Höfer, Weiner and Feldon[69] | Healthy adults: smokers and non-smokers | Greater trials to criteria in PE compared with no-PE group. | Smoking: main effect not reported. | No main effect or interactions<br>**No sex difference in LI, or interaction with smoking status or schizotypy.** |
|  |  |  | Both visual and auditory: greater trials to criteria in PE compared with no-PE group. | Schizotypy/smoking: main effects not reported. | No main effect or interactions<br>**No sex difference in LI, or interaction with smoking status or schizotypy.** |
| 1999 | Höfer, De la Casa and Feldon[70] | Healthy adults | No main effect of PE condition. | Schizotypy: main effect not reported. | No main effect or interactions<br>**No sex difference in LI, or interaction with schizotypy.** |
| 2001 | Lubow, Kaplan and De la Casa[64] | Adults (university students) | Reduced reaction time in PE compared with no-PE group. | Schizotypy: no main effect. | Sex: significant effect (direction not reported).<br>Near-significant sex X pre-exposure X schizotypy interaction ($p = 0.065$).<br>Main effect of PE in males but no PE X schizotypy interaction.<br>Main effect of PE in females with PE X schizotypy interaction: reduced LI in high schizotypal compared with low schizotypal females. |
|  |  |  |  |  | **Reduction of LI with increased schizotypy in females but not males.** |
| 2001 | Wuthrich and Bates[71] | Adults (university students and others) | Greater trials to criteria in PE compared with no-PE group. | Schizotypy: main effect not reported. | No main effect or interactions<br>**No sex difference in LI, or interaction with schizotypy.** |
| 2002 | Lubow and De la Casa[72] | Adults (university students) | Greater number of correct predictions in PE compared with no-PE group (marginal effect, $p = 0.07$). | Schizotypy: no main effect. | No main effect of sex.<br>Interaction between sex and schizotypy: LI observed in high-schizotypal male and low schizotypal female participants. No LI in low-schizotypal male and high schizotypal female participants.<br>**Opposing effects of schizotypy on LI between males and females.** |
| 2007 | Evans, Gray and Snowden[73] | Healthy adults | Increased reaction time in PE compared with no-PE group.<br>Increased correct responses in no-PE compared with PE group. | Greater LI (indexed by correct responses) in individuals with higher impulsive non-conformity (no effect on reaction time).<br>Reduced LI (indexed by correct responses) in individuals who smoke with high unusual experiences (no effect on reaction time). | No main effect or interactions.<br>**No sex difference in LI, or interaction with smoking status or schizotypy.** |
| 2009 | Shrira and Kaplan[74] | Adults (university students) | Increased correct responses in no-PE compared with PE group. | LI observed in low interpersonal schizotypy factor but not in high interpersonal schizotypy factor. | No main effect of sex.<br>Interaction between sex and cognitive-perceptual schizotypy sub-factor: low cognitive-perceptual male and high cognitive-perceptual female participants<br>showed LI, whereas high cognitive-perceptual male and low cognitive-perceptual female participants did not show LI.<br>**Opposing effects of cognitive-perceptual schizotypy factor on LI between males and females.** |
| 2009 | Shrira and Tsakanikos[75] | Healthy adults (university students) | Increased correct responses in no-PE compared with PE group. | Increased unusual experiences schizotypy factor associated with lower LI.<br>Increased introverted anhedonia (negative schizotypy) factor associated with greater LI. | No main effect or interactions.<br>**No sex difference in LI, or interaction with schizotypy.** |
| 2010 | Lee and Telch[76] | Adults (university students) screened for OCD symptoms (highest and lowest scorers retained) | Greater reaction time in PE compared with no-PE condition for control and reactive obsession OCD, but not in autogenous obsession OCD. | No main effect of schizotypy on LI. | **Main effect: Greater LI in female compared with male participants.**<br>No interaction between sex and obsessive-compulsive symptoms or schizotypy. |
| 2011 | Kaplan and Lubow[77] | Healthy adults (university students) | Greater reaction time in PE compared with no-PE group. | No main effect of schizotypy. | No main effect of sex.<br>Interaction between sex and schizotypy on LI score. No LI observed in low schizotypal female and high schizotypal male participants. LI observed in high schizotypal female and low schizotypal male participants.<br>**Opposing effects of schizotypy on LI**<br>**between males and females.** |
| 2012 | Granger, Prados and Young[78] | Adults (university students) | Greater reaction time in PE compared with no-PE group. | Increased unusual experiences factor of schizotypy associated with reduced LI score. | No main effect or interactions.<br>**No sex difference in LI, or interaction with schizotypy.** |
| 2022 | Dawes, Quinn, Bickerdike... Waddington, O'Tuathaigh and Moran[79] | Healthy adults (university students) | Greater reaction time in PE compared with no-PE group. | Aberrant salience: higher Aberrant Salience Inventory score associated with reduced LI.<br>Schizotypy/cannabis use: no main effects on LI score. | No main effect of sex (interactions not tested).<br>**No sex difference in LI.** |
| **Studies in schizophrenia patients** |  |  |  |  |  |
| 1996 | Swerdlow, Braff, Hartston, Perry and Geyer[80] | Schizophrenia/OC D/affective disorder patients and healthy controls | Auditory and Visual task:<br>Greater trials to criteria in PE compared with no-PE group. | Auditory and visual study:<br>No effect of schizophrenia patient status or smoking on LI.<br>Auditory only:<br>No effect of age, psychosis proneness or affective disorder/OCD patient status on LI. | Auditory and visual study:<br>No main effect of sex. Not stated whether effect of sex on LI was assessed in schizophrenia patients.<br>**No sex difference in LI.** |
| 2000 | Lubow, Kaplan, Abramovich, Rudnick and Laor[81] | Adult schizophrenia patients and controls | Greater reaction time in PE compared with no-PE group. | No significant two-way interaction between patient status and LI condition. | Interaction between patient status and sex: LI present in male control participants and schizophrenia patient participants of both sexes. In female schizophrenia patients, LI reversal observed with faster responding in PE compared with no-PE group.<br>**Sex difference in effect of schizophrenia patient status on LI.** |
| 2001 | Serra, Jones, Toone and Gray[65] | Schizophrenia patients and their | No main effect of pre-exposure condition. | Clinical group: no main effect or interaction with PE condition. | No main effect or interactions.<br>**No sex difference in LI.** |
|  |  | schizotypal and non-schizotypal relatives |  |  | (Not stated whether interaction between sex and schizophrenia patient status was assessed). |
| 2002 | Leumann, Feldon, Vollenweider and Ludewig[82] | Adult schizophrenia patients (atypical vs typical antipsychotics) and controls | Greater trials to criteria in PE compared with no-PE group. | No interaction between pre-exposure group and patient status/medication condition. | No main effect of sex. Not stated whether interactions were tested.<br>**No sex difference in LI.** |
| 2005 | Swerdlow, Stephany, Wasserman ... Sharp, Minassian and Auerbach[83] | Adult schizophrenia patients and controls | Greater number of correct trials in no-PE compared with PE group. | Lower LI in high harm avoidance individuals.<br>Significant LI observed in low-smokers but no LI observed in high smokers.<br>No interaction between pre-exposure and other personality measures (Tri-dimensional Personality Questionnaire/Sensation Seeking Scale) or age.<br>No effect of schizophrenia patient status on LI. | No main effect of sex or interaction with PE condition or personality measures.<br>Three-way interactions between sex, personality measures and LI not reported. However, when men and women were split, negative association between LI and harm avoidance only seen in women. Association did not differ between follicular and luteal phase.<br>In women, no effect of menstrual phase on LI. Not stated whether interaction between sex and schizophrenia patient status was assessed.<br>**Female-specific association between harm avoidance and LI.** |
| 2009 | Gal, Barnea, Biran ... Feldon, Fennig and Levkovitz[84] | Adult schizophrenia patients and matched controls | Greater reaction time in PE compared with no-PE groups. | LI only observed in controls but not schizophrenia patients during trials 1-5. LI observed in both groups during trials 6-10. | Tendency towards greater LI in female participants ($p = 0.07$) during trials 1-5.<br>No interaction between sex and schizophrenia patient status. |
|  |  |  |  |  | **Greater LI in females compared with males.** |
| **Studies in other clinical cohorts** |  |  |  |  |  |
| 1993 | Lubow and Josman[85] | Hyperactive and typically developing children (5-7 years old). | Hyperactive and control participants analysed separately.<br>No effect of PE on trials to criteria in either control or hyperactive group. | Interaction between age and pre-exposure in controls. Post hoc tests show LI observed in younger children but reversal of LI in older children in control participants.<br>No interaction between age and pre-exposure in hyperactive participants. | Effect of sex not assessed in hyperactive children owing to too few females. Analyses in hyperactive group conducted both with and without females for sensitivity and results did not differ.<br>No main effect of sex on LI in controls or interaction with age.<br>**No sex differences in LI.** |
| 1999 | Lubow, Dressler and Kaplan[86] | Adult Parkinson's Disease patients and controls | Greater reaction time in PE compared with no-PE group. | No interaction between PE and PD patient status. | No sex difference in LI or interaction with PD patient status.<br>In PD patients, significant three-way interaction between PE, sex, and PD symptom side: LI only observed in left-sided males and right-sided females, and LI was reversed in left-sided PD females.<br>**Sex difference in effect of PD symptom side on LI.** |
| 1999 | Swerdlow, Hartston and Hartman[87] | Obsessive compulsive disorder (OCD) patients and healthy controls | Greater trials to criteria in PE compared with no-PE group. | Greater LI effect in OCD patients compared with healthy controls. | No effect of sex on LI. Not stated whether interaction between sex and schizophrenia patient status was assessed.<br>**No sex difference in LI.** |
| 2000 | Lubow, Toren, Laor and Kaplan[88] | Children and adolescents; | Greater reaction time in PE compared with no-PE | No interaction between PE group and anxiety level. | No sex difference in LI or interaction with anxiety disorder status. |
|  |  | anxiety disorder diagnosed and controls | group. |  | **No sex differences in LI.** |
| 2017 | Györfi, Nagy, Bokor, Kelemen and Kéri[89] | Adult Parkinson's Disease patients and matched controls | Greater reaction time in PE compared with no-PE group. | Stronger LI in non-medicated PD patients compared with controls. Weaker LI in medicated (L-DOPA) PD patients compared with controls. | No sex difference in LI in PD patients (not reported in controls).<br>**No sex difference in LI.** |

LI is demonstrated if the participant is faster to respond to the non-pre-exposed cue compared with the pre-exposed cue (reaction time tasks) or takes fewer trials to learning criteria in the non-pre-exposed compared to pre-exposed condition (trials to learning criteria tasks). Auditory designs use the same principles, but cues are typically white noise tones and targets are typically nonsense syllables.

The majority of experiments (20/31 from 17 separate studies) assessed LI in non-clinical cohorts, with or without assessing the effect of schizotypy on LI. In the 6 non-clinical experiments that did not assess the impact of schizotypy (Table 1: Studies in non-clinical adult cohorts), no sex differences in LI were found in any study. However, when the effect of schizotypy was considered (Table 1: Studies assessing schizotypy in non-clinical adult cohorts), results are inconsistent. Nine experiments across eight studies did not show sex differences in the effects of schizotypy on LI or main effects of sex on LI. Two studies in the same group showed similar findings of stronger LI in low schizotypal compared with high schizotypal females[64,72]. However, inconsistent effects in males were observed between the two studies with no LI observed in males and no interaction with schizotypy in the earlier study, but the opposite effect to that observed in females found in the later study. A later study by the same group once again found sex differences. However, the effect in females was reversed to what had been seen in the earlier studies, with LI only observed in high-schizotypal but not low-schizotypal participants, with the opposite trend observed in males[77]. Shrira and Kaplan (2009)[74] assessed the effect of specific schizotypy dimensions and found an interaction between sex and schizotypy specifically in the cognitive-perceptual dimension, with males scoring high on this dimension and females scoring low showing attenuated LI. Lee and Telch (2010)[76] did not observe an interaction between sex and schizotypy, but did find a main effect of sex, with greater LI demonstrated in female compared with male participants, contrasting the findings of the studies in non-clinical cohorts that did not assess schizotypy. Overall, no evidence for sex differences was found in non-clinical cohorts that did not assess schizotypy. However, in studies assessing the effect of schizotypy in non-clinical cohorts, there was mixed evidence for an effect of sex.

Sex differences in LI in schizophrenia cohorts were often unclear, as studies typically report only main effects of sex rather than its interactions with patient/control status or medication status. Hence, it cannot be discerned from these studies whether LI disruption in schizophrenia is itself sex-dependent. Notwithstanding, no overall sex effects were seen in half of the studies (3/6). However, one study showed reversal of LI learning (i.e., greater learning performance in the pre-exposed compared to non-pre-exposed condition) in female schizophrenia patients[81], and another did not find an interaction between sex and schizophrenia patient/control status but did show a tendency towards stronger LI in females compared with males across all participants[84]. Swerdlow et al. (2005)[83] did not find that sex interacted with schizophrenia patient status, rather harm avoidance was measured as a personality variable in participants, and it was found that there was a negative association between harm avoidance and LI in women, that was not seen in men. Unfortunately, it was not assessed whether this interacted with schizophrenia patient status. Five studies in other clinical cohorts were found, the majority of which did not show sex differences in LI[81,85,87,89]. However, Lubow et al. (1999)[86] found that the sidedness (left or right) of symptoms in Parkinson’s disease had a different association with LI between men and women, with LI only observed in left-sided males and right-sided females, and LI was reversed in left-sided females. Similarly to the non-clinical cohorts, there was mixed evidence for an effect of sex in schizophrenia cohorts, but minimal evidence for an effect of sex in studies in other clinical cohorts. Taken together, sex does not appear to reliably influence LI when assessed in isolation but exerts inconsistent effects when included as a variable alongside schizotypy or clinical status.

#### 4.2.3. Review of rodent model studies

Although latent inhibition has been extensively studied in rodents for several decades, only a minority of studies have used both male and female animals, and only a subset of those have also included sex as a factor in statistical analyses or alternatively stratified their analyses by sex. In this review, a total of 60 experiments across 54 studies that met the inclusion criteria were identified (Table 2).

**Table 2.**
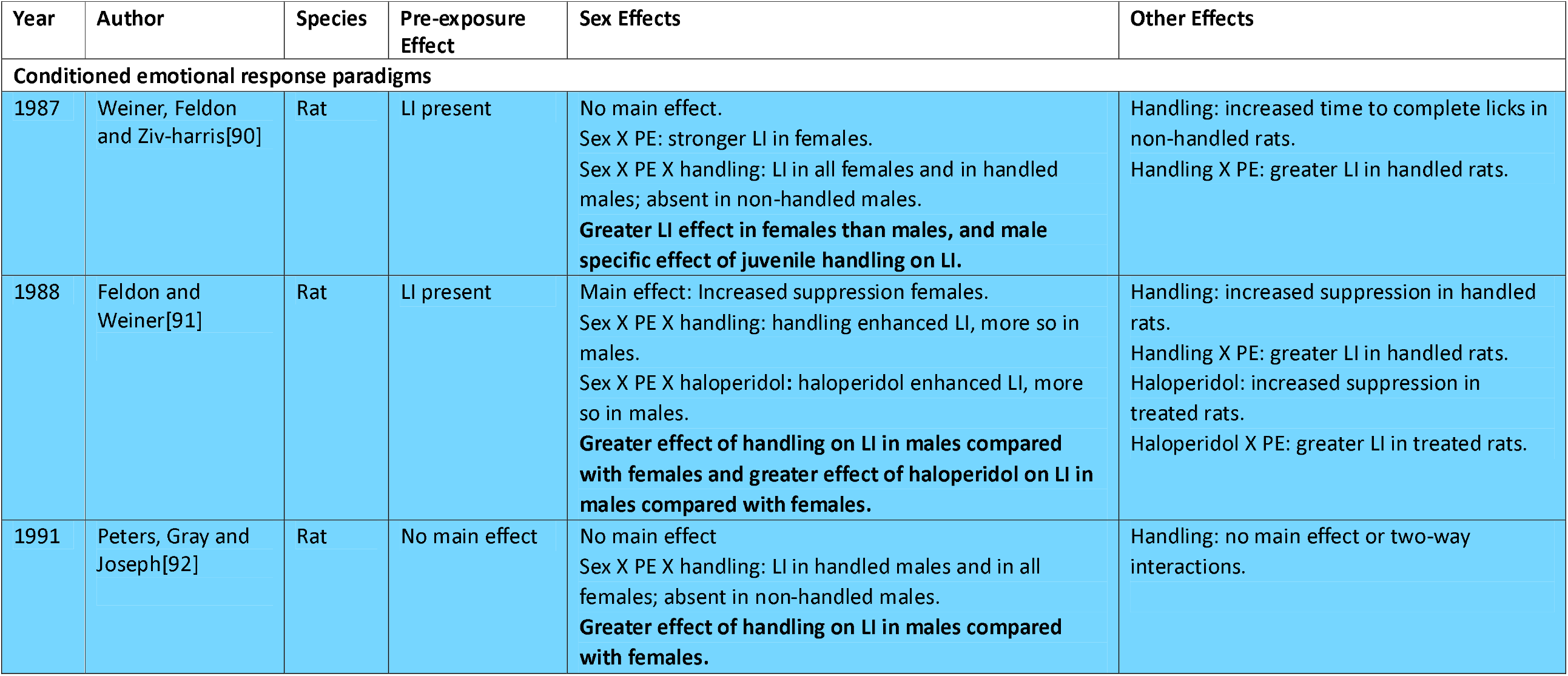
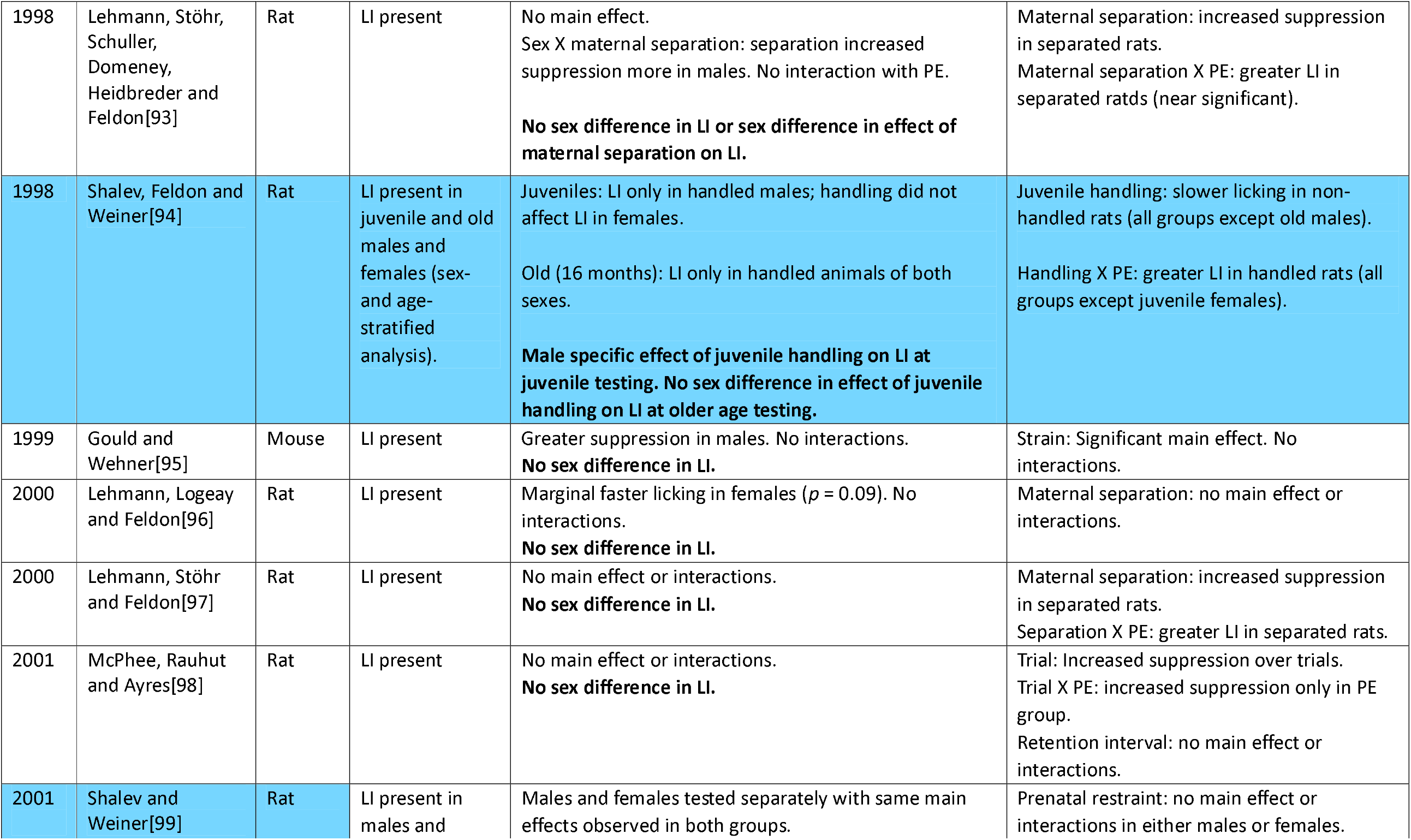
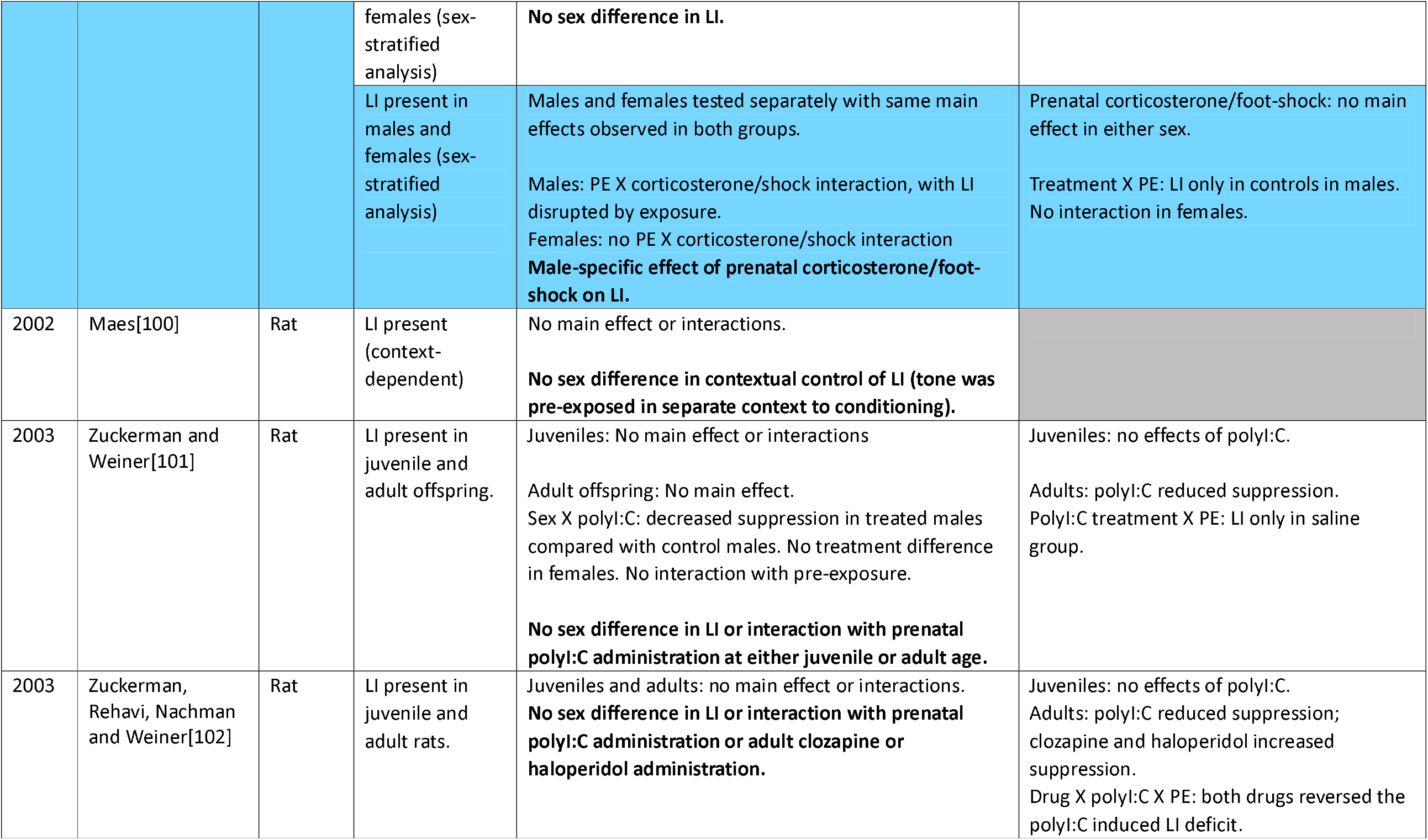
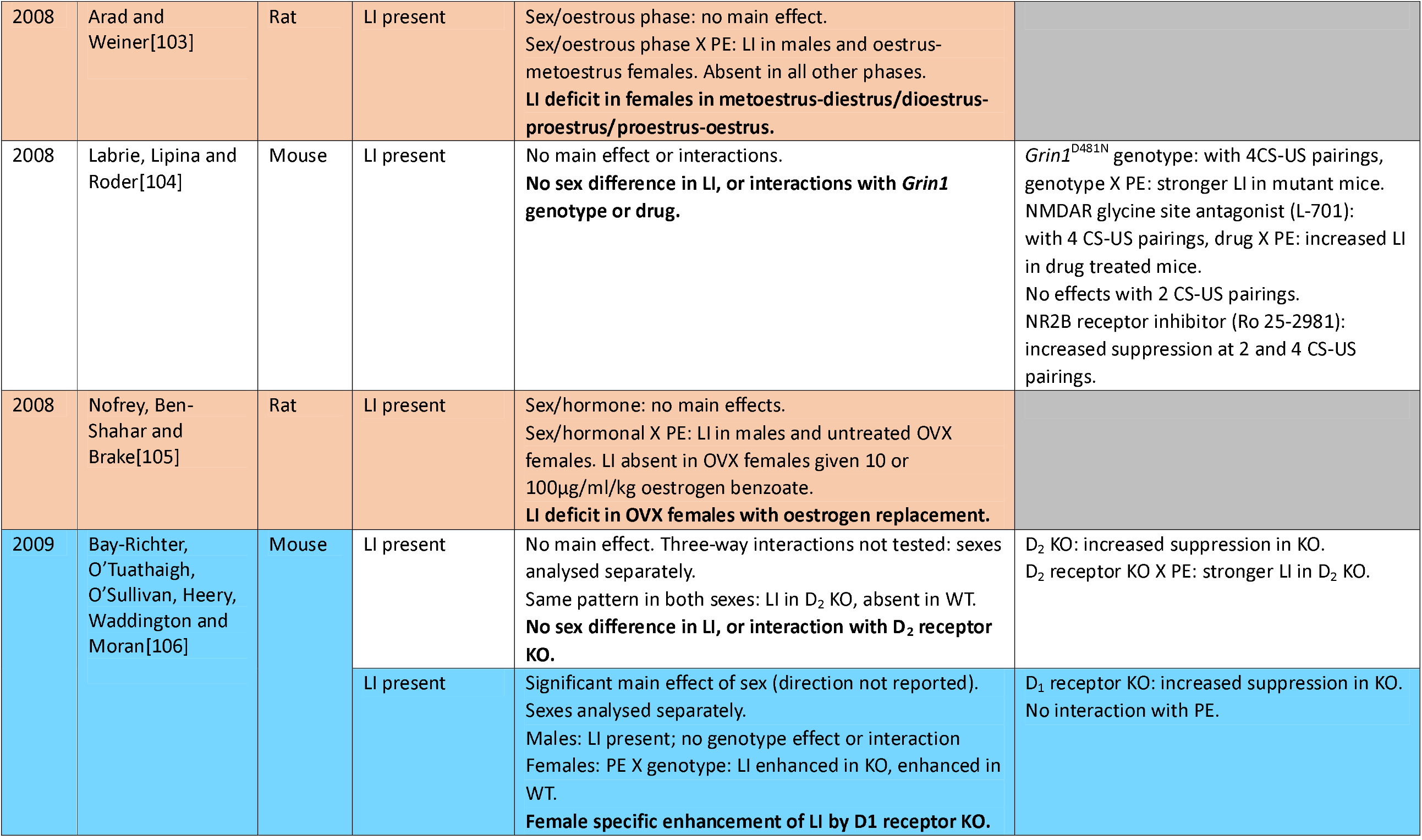
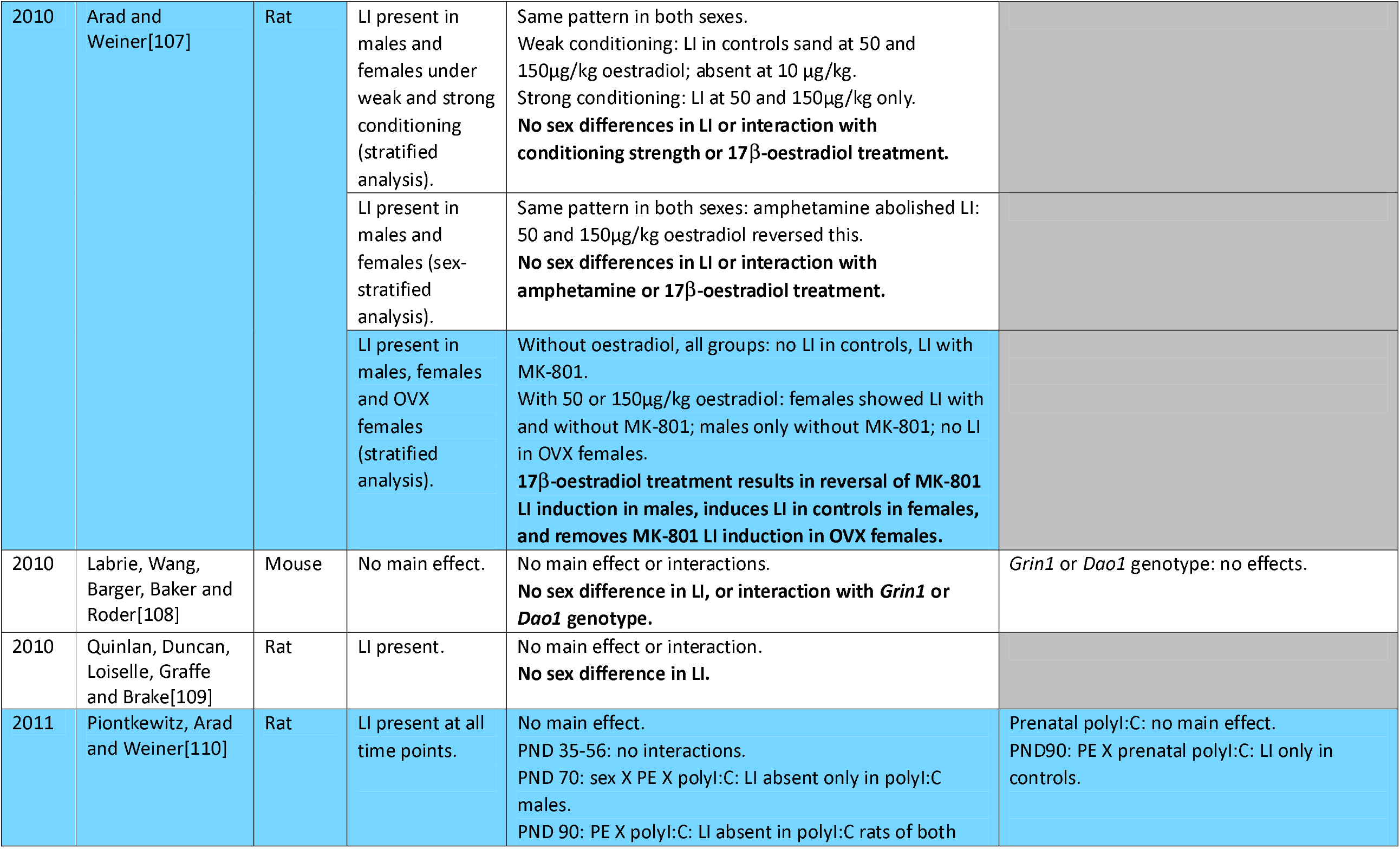
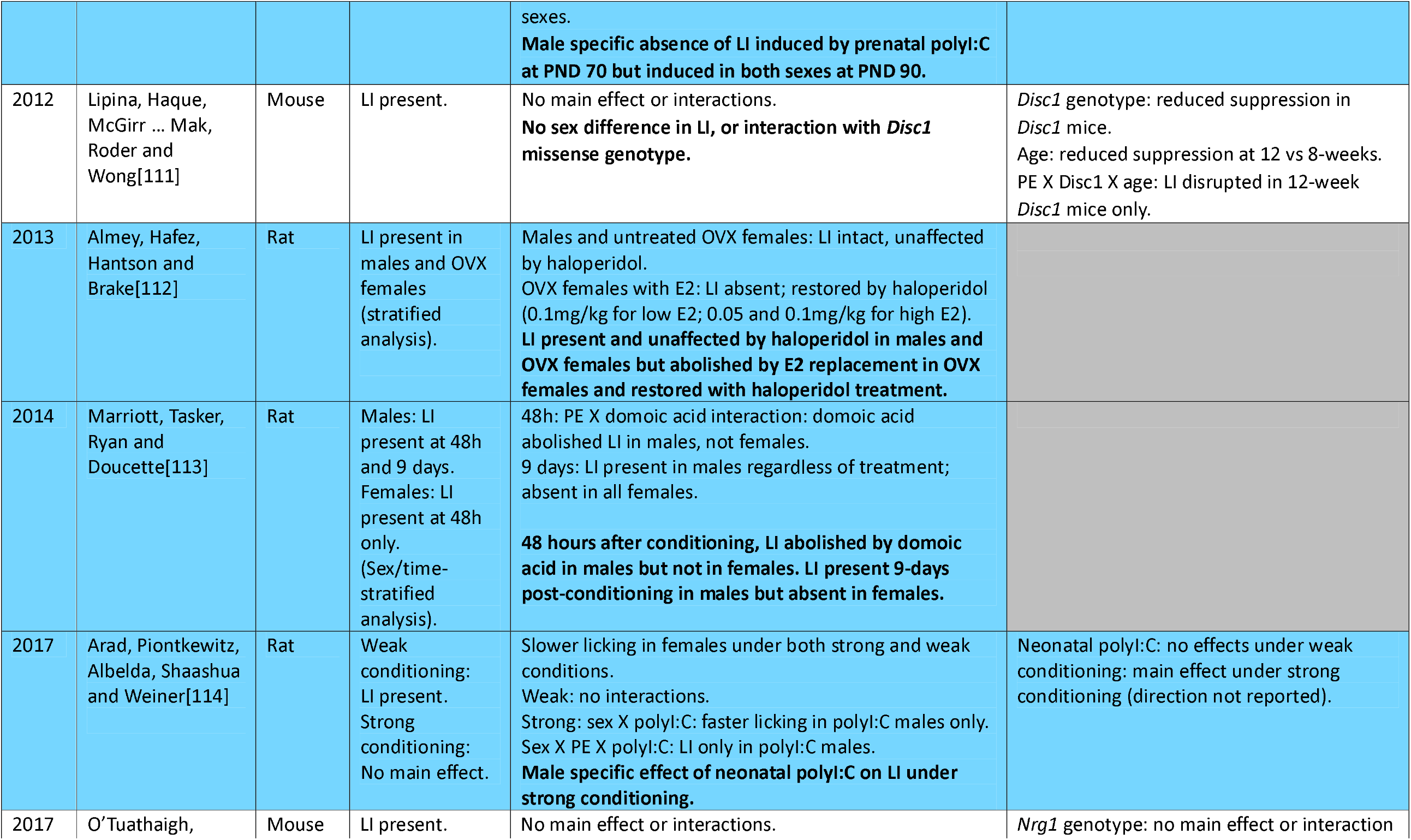
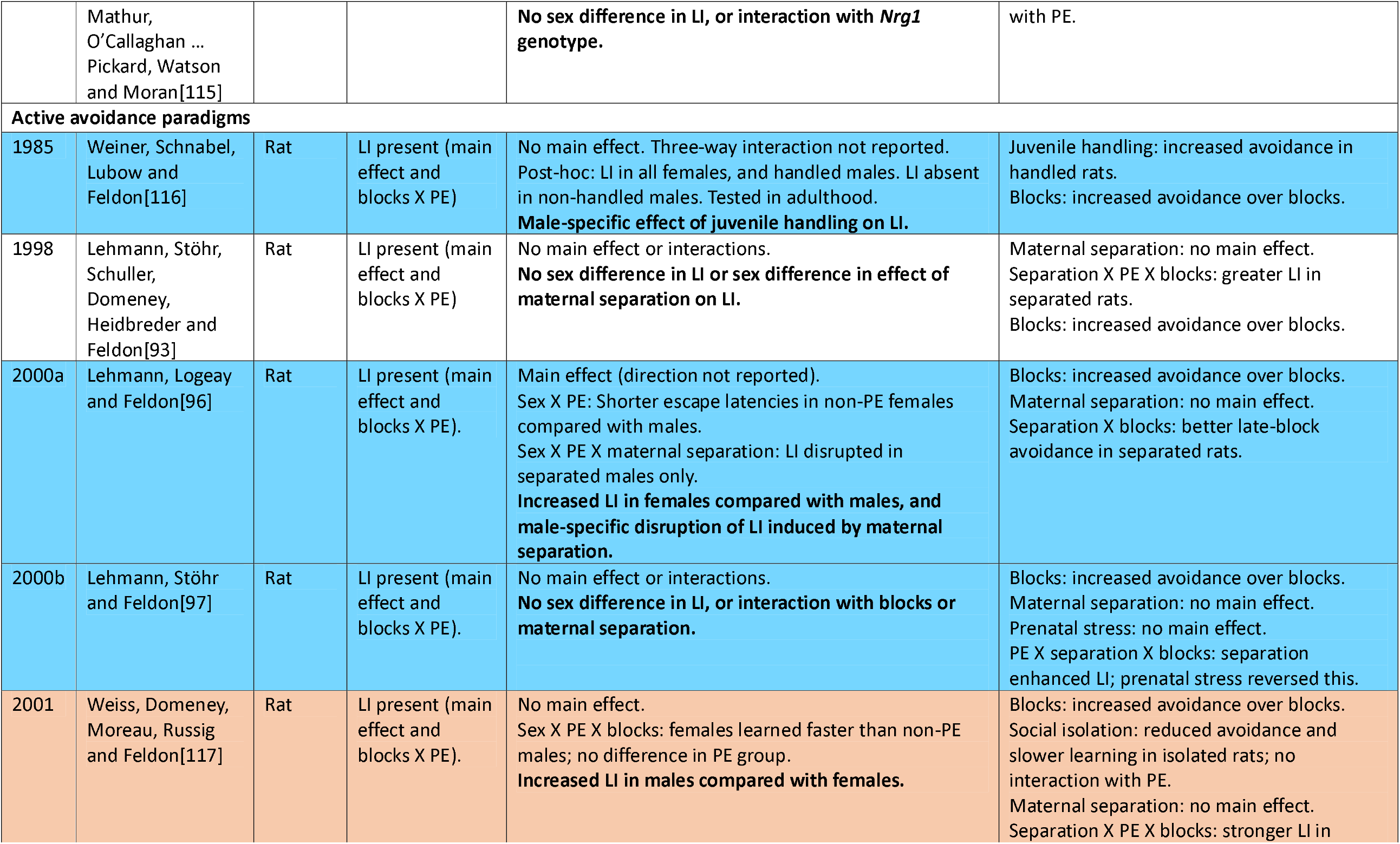
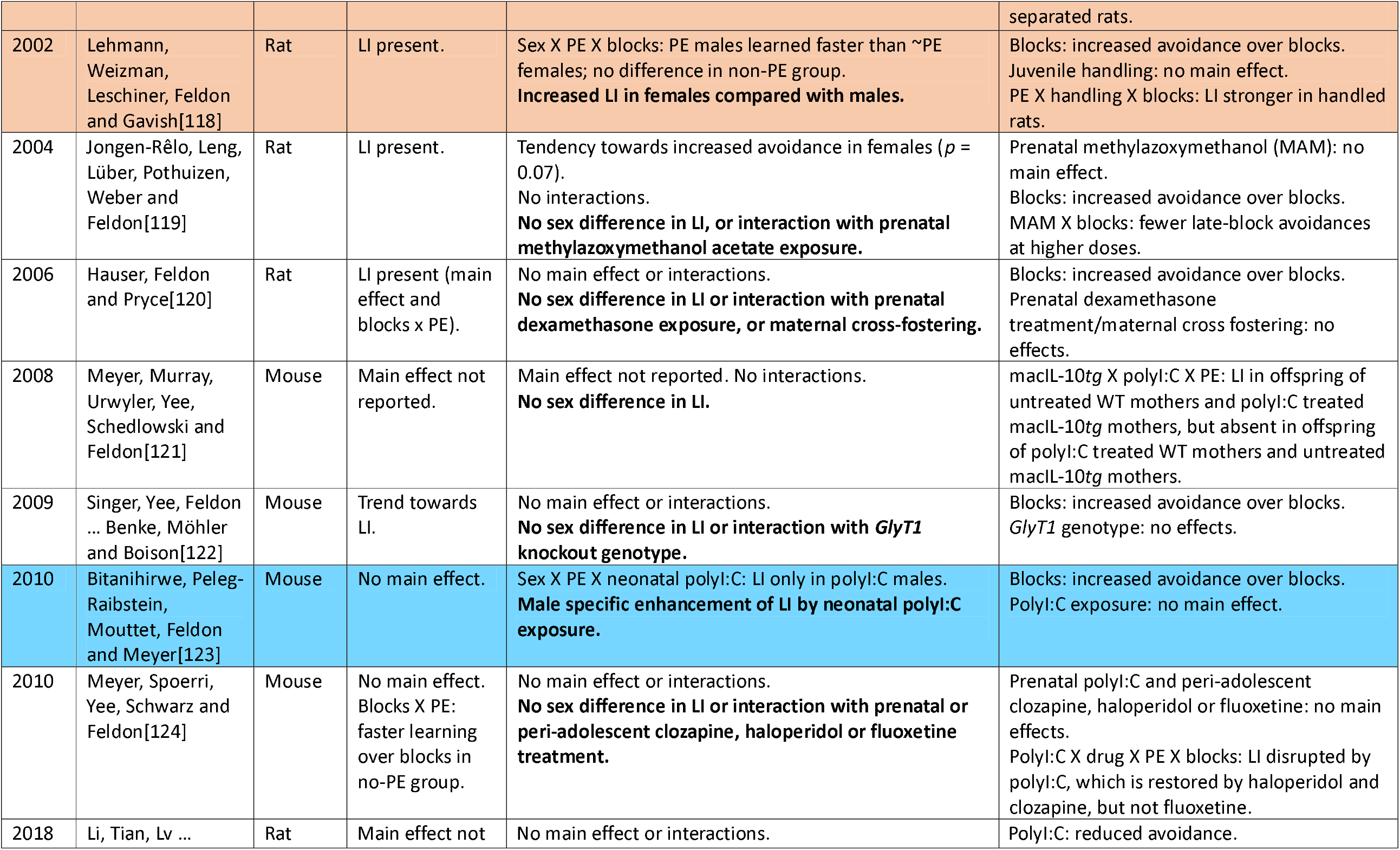
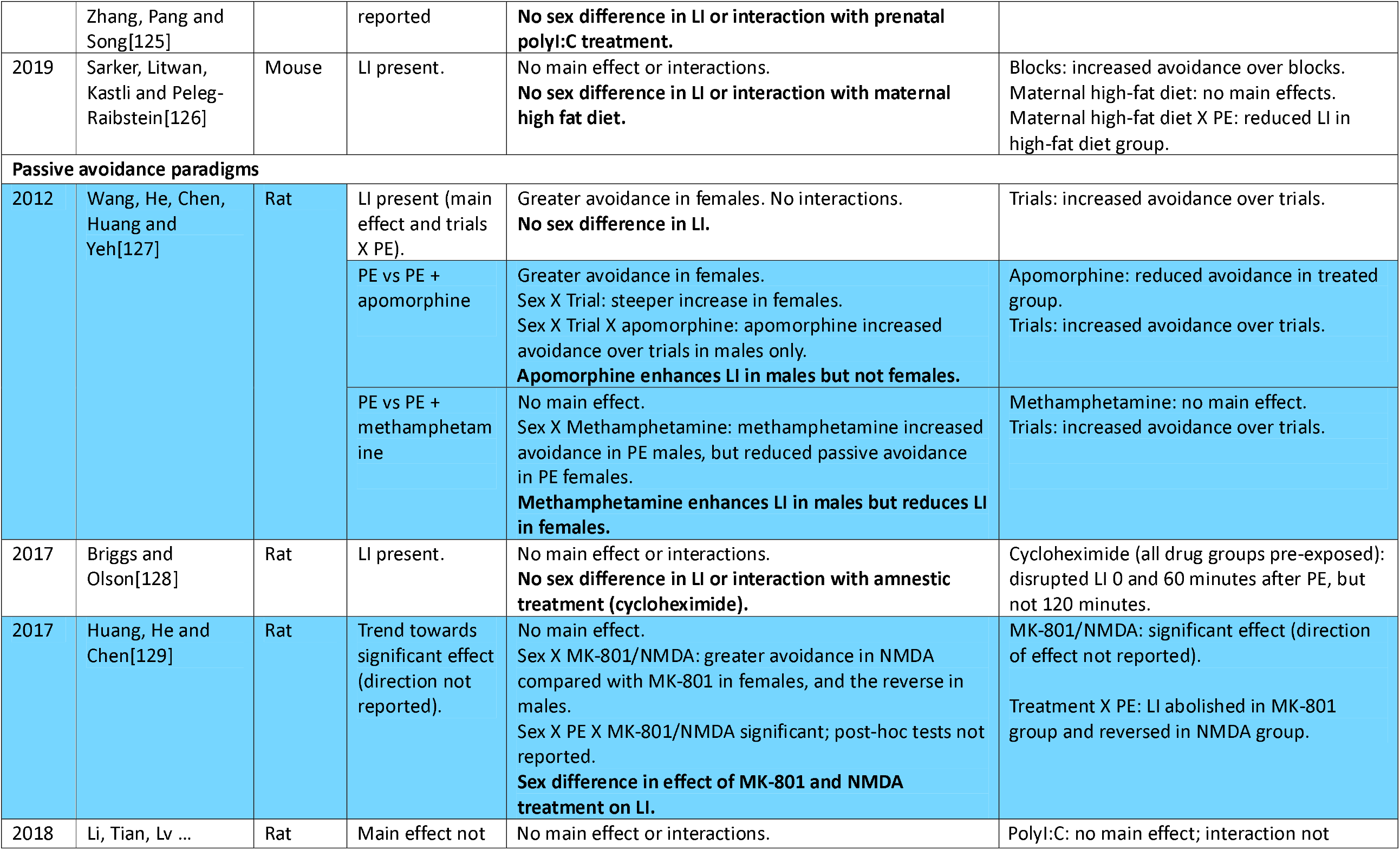
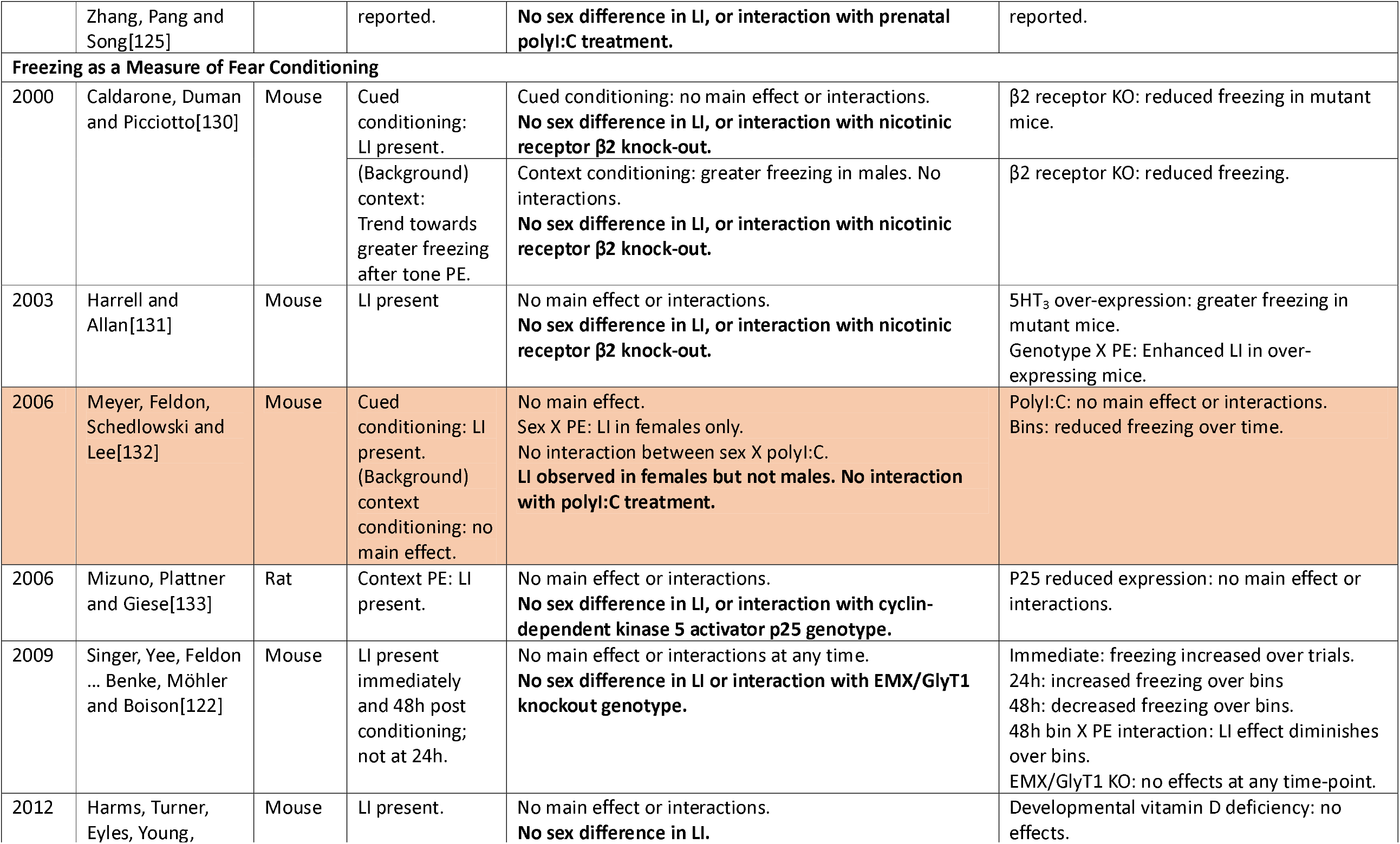
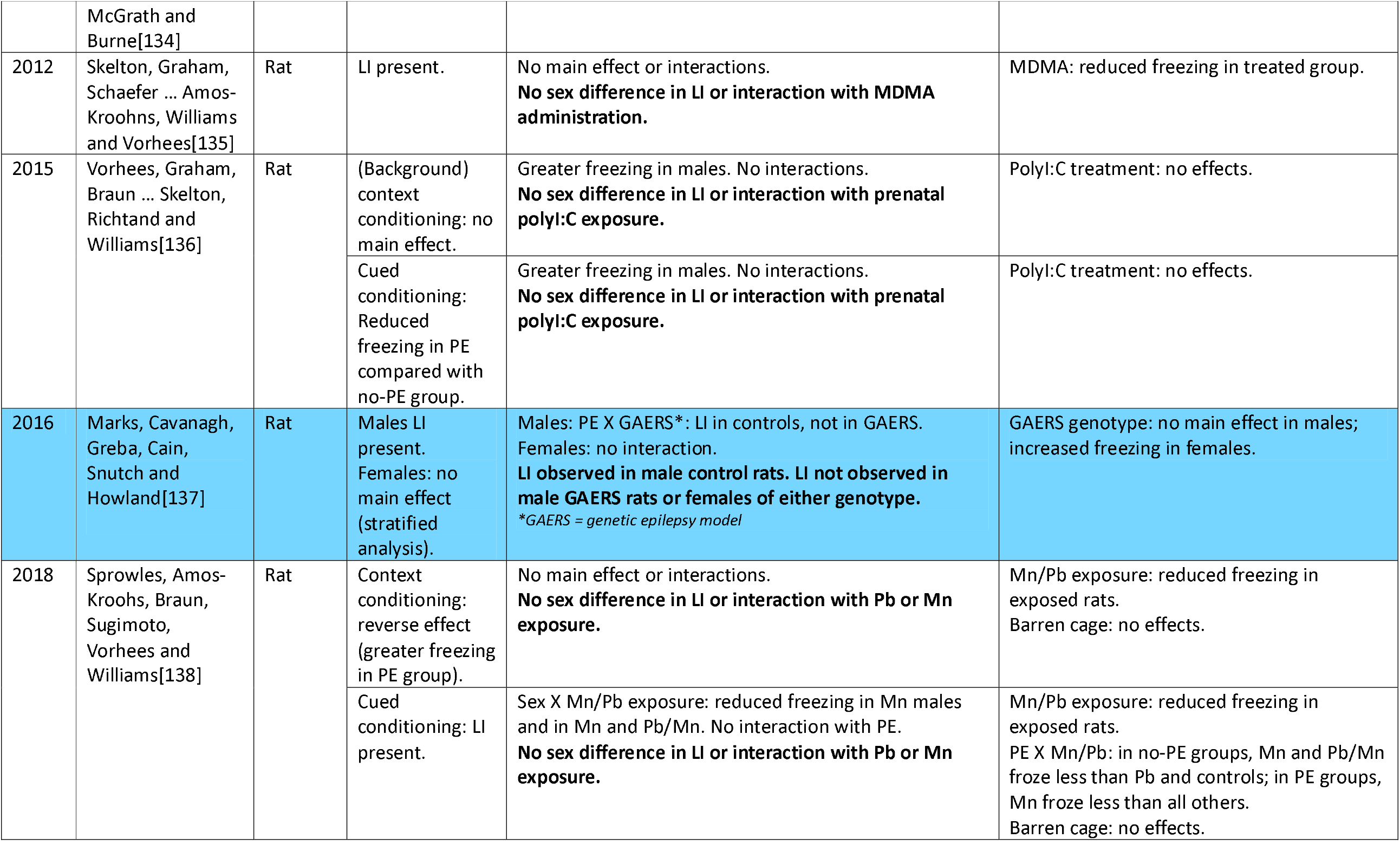
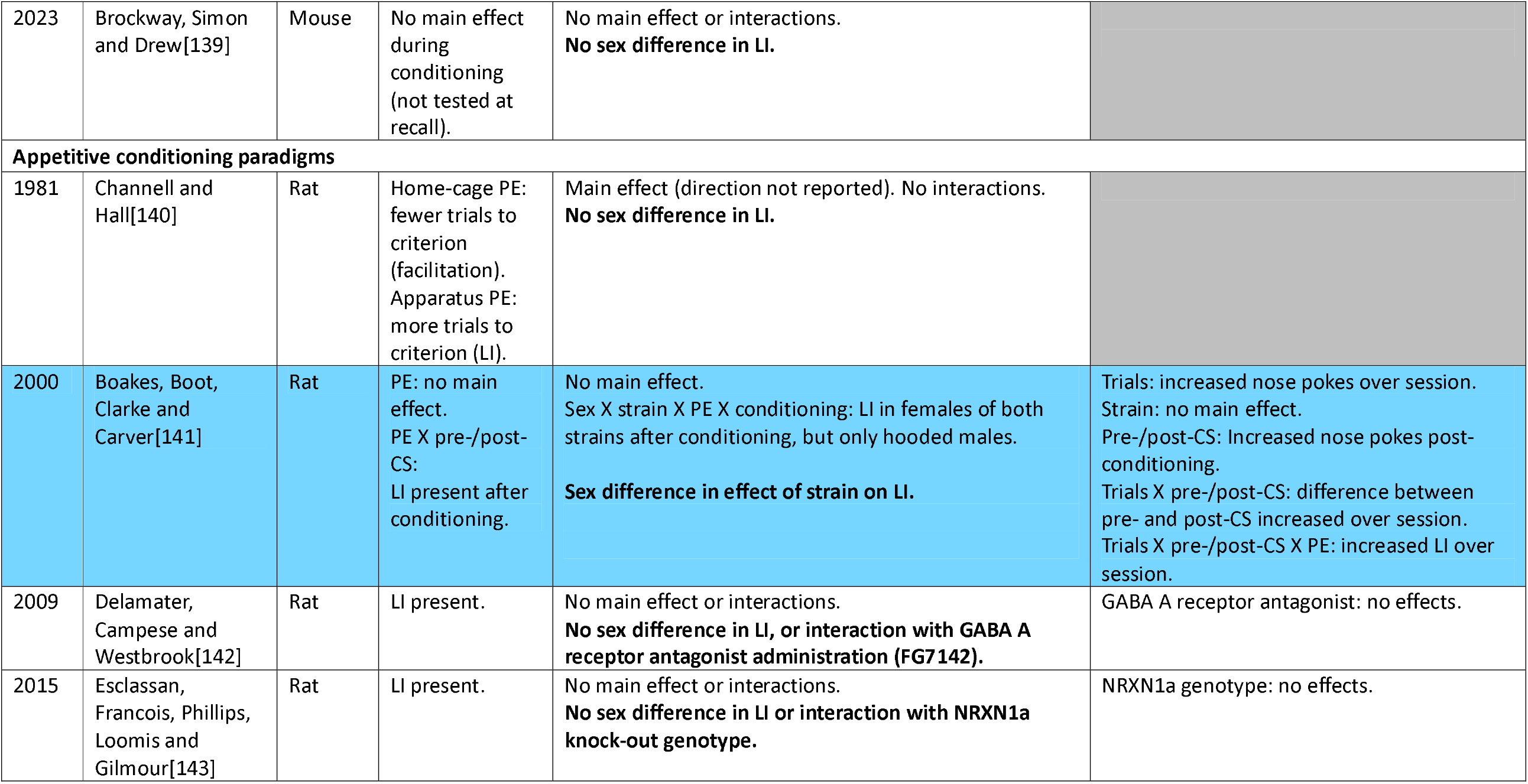
Sex differences in latent inhibition in rodents. Studies highlighted in orange show an overall effect of sex on LI (interaction between sex and pre-exposure). Studies highlighted in blue show an effect of sex on the relationship between LI and an additional variable (three-way interaction between sex, pre-exposure, and additional variable). Conclusions relating to sex differences in LI in each study are highlighted in bold. OVX: ovariectomisation; PE: pre-exposure.

A range of behavioural paradigms were used across LI studies in rodents, with the majority using an aversive US, primarily foot-shock (55/59), and a discrete stimulus cue (tone/light), rather than the context only, acting as the CS (51/59). The most common cued paradigm was CER (26/59), in which the suppression of drinking in water-restricted animals in response to the CS serves as the conditioned response. Fourteen experiments used active avoidance, with escape latency to a separate compartment as the measure of conditioning, and four used passive avoidance, assessing only contextual conditioning via latency to re-enter the conditioning compartment. Eleven experiments used freezing as a measure of fear conditioning (six cued, one solely contextual, and four both), with total freezing time at recall as the dependent measure. In equivalent contextual conditioning studies, contextual fear conditioning can either be foreground or background[4]. In background CFC, the US is paired with a discrete cue CS, but CFC is assessed by measuring time spent freezing when the animal is returned to the context in which conditioning occurred. Conversely, in foreground CFC, there is no discrete cue CS, and the context is the only predictor of the US. Four studies used appetitive rather than fear based conditioning models in which the animal develops an association between the CS and a food-based US, all of which assessed cued conditioning.

The majority of studies did not find sex effects in LI (37/59 experiments). Sex differences were found in wild-type animals at baseline in a minority of experiments, three of which showed stronger LI in females compared with males[96,118,132] whereas two studies showed stronger LI in males compared with females[117,137]. However, in numerous instances, sex was found to interact with an additional variable to affect LI, such as genotype, juvenile handling or drug administration. Nine CER experiments demonstrated interactions with sex across juvenile handling[90–92,94], corticosterone[99], domoic acid[113], and polyI:C administration, and D1 receptor genotype[106]. Such interactions were also found in passive avoidance studies, with sex differences seen in the effects of polyI:C administration[123] and juvenile handling[116] on LI. Interactions between drug administration and sex on LI were seen by Wang et al. (2012)[127] in the case of ascorbic acid, methamphetamine, but not apomorphine. In passive avoidance models, Huang et al. (2017)[129] assessed the impact of NMDA and MK-801 administration on LI and reported an interaction between sex and drug administration on LI, however specific post hoc comparisons were not reported. In contrast, using fear conditioning Meyer et al. (2006)[132] did not find an interaction between sex and polyI:C treatment of dams.

Five studies (all CER paradigms) assessed the effect of oestrous phase or ovariectomisation and oestrogen replacement in females on LI, which reported effects of oestrogen or ovarian hormones on LI[103,105,107,109,112]. The rodent oestrus cycle consists of four phases: dioestrus, proestrus, oestrus and metoestrus, with ovulation occurring between proestrus and oestrus phases, and oestrogen peaking during proestrus[144]. In studies that assessed LI with female rats stratified across the oestrous cycle, in all cases LI was disrupted when conditioning took place during the proestrus phase[103,107,109]. Similarly, ovariectomised rats receiving oestrogen replacement showed attenuated LI whereas vehicle treated ovariectomised rats displayed normal LI[105]. This evidence suggests that variation in ovarian hormones may contribute to sex differences in LI and could obscure sex effects when hormonal status is not considered. Fully characterising the influence of oestrogen on LI is of particular relevance to psychosis, as a later life incidence of schizophrenia in women coincides with menopausal transition[145]. Furthermore, the effect of oestrous replacement in ovariectomised females was found to interact with administration of the antipsychotic haloperidol[112], raising further implications for sex differences in mechanisms underpinning psychosis.

Most studies included in the review that assessed the effect of sex on LI under baseline conditions did not observe sex differences in LI. However, several studies identified interactions between sex and additional factors such as drug administration, juvenile handling and circulating levels of oestrogen in females. These findings suggest sex may influence the neurobiological substrates underlying LI, despite similar performance between males and females under baseline conditions, with sex-dependent effects emerging under specific experimental manipulations[146]. Human studies offered limited evidence for sex differences in LI. Hence, the complex interactions between LI, sex, and additional variables in rodents highlight the value of animal model studies for probing the underlying biological mechanisms that may contribute to a sexually differentiated vulnerability to disruptions in LI, even when performance under baseline conditions is similar between males and females.

## 5. Sex differences in the relationship between PE duration and LI of contextual fear memory

The systematic review found limited evidence for sex differences in LI at baseline in both humans and rodents. However, several rodent model studies assessed the impact of additional factors on LI such as drug administration, juvenile handling, and oestrous phase, and in many instances, interactions between sex and these factors were observed. Hence, although LI performance may be similar in males and females at baseline, there is evidence that sex may influence the effects of various experimental manipulations on LI. Consequently, given the relevance of LI to cognitive processing implicated in psychosis, it is important to understand the mechanisms for this sexually differentiated vulnerability of LI to disruption.

The formation of configural representations of contexts is mediated by the hippocampus[30], and the hippocampus is also crucial for the formation and retrieval of associations between context and CS and outcomes, resulting in the context-dependency of cued fear conditioning and LI[11,147]. Foundational studies on sex differences in behavioural, molecular and circuit mechanisms of memory have used foreground CFC to show that the optimal training conditions and the molecular and circuit mechanisms supporting learning may differ between males and females[148]. Hence, here we have further explored sex differences in both the behavioural and hippocampal molecular underpinnings of LI using a CFC paradigm.

We compared LI of CFC in male and female Sprague-Dawley rats across a range of context pre-exposure (PE) durations (5, 20 or 60 minutes per day for three consecutive days, Fig.2A). ARRIVE guidelines[149] were adhered to, and ethical approval was obtained from the Cardiff University Animal Welfare and Ethical Review Body.

**Fig. 2.**
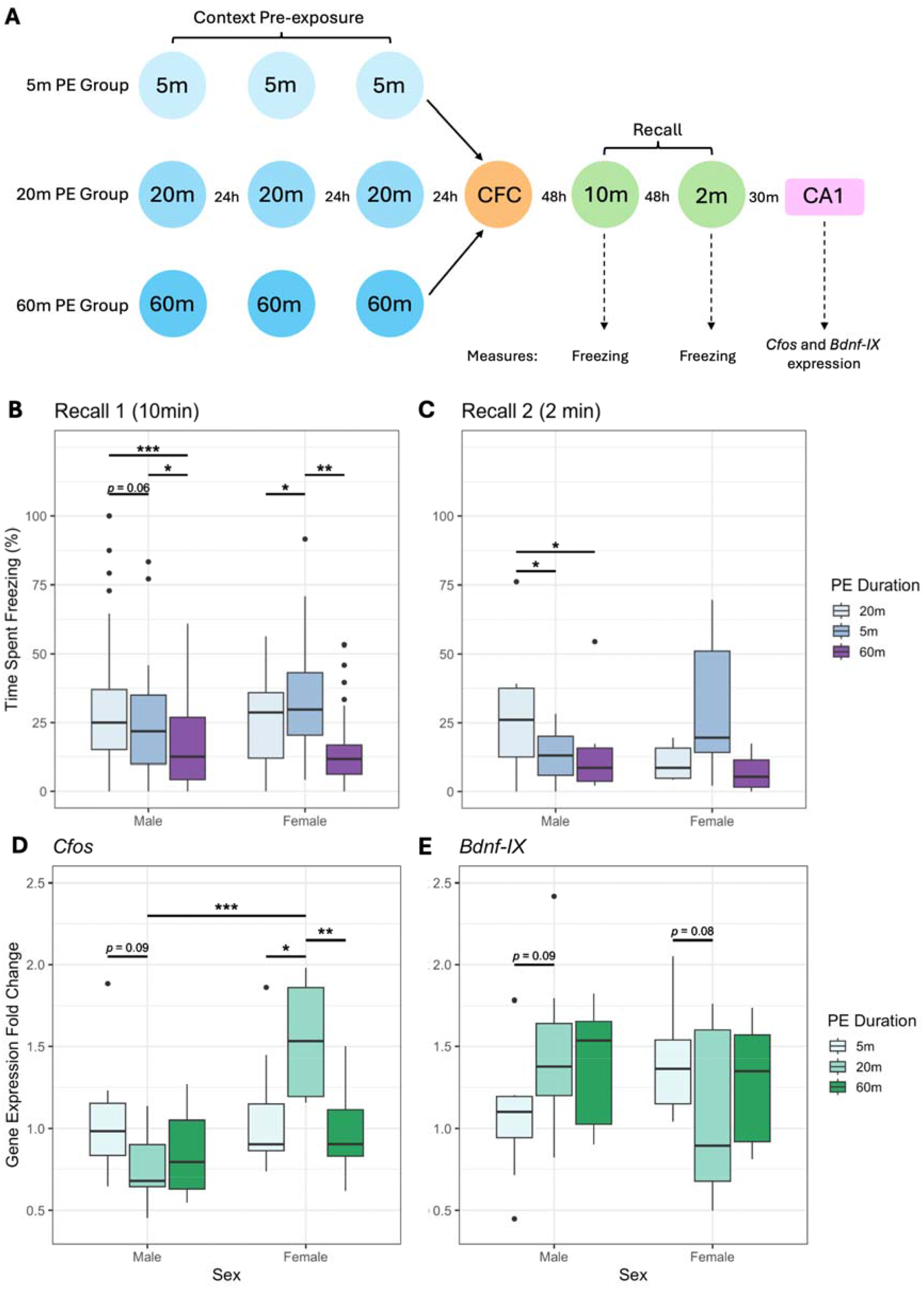
**A**. Schematic of pre-exposure (PE), contextual fear conditioning (CFC) and recall protocol. Rats were pre-exposed to the conditioning context for either 5, 20 or 60min per day over 3d. CFC was established 24h later using a 2s 0.64mA scrambled foot-shock, given 2min into a 3min exposure to the context. Rats were returned to the context for a 10min recall trial 2d later and a 2min recall trial a further 2d later. Rats were euthanised (rising concentration of CO_2_) 30min after the final recall trial and CA1 tissue was collected for qPCR analysis of *Cfos* and *Bdnf-IX* expression. All trials were run at the same time each day for each rat. Two-way ANOVAs showed interactions between Sex and PE Duration for freezing behaviour during both the ten-minute (**B**) and two-minute (**C**) recall sessions, and in *Cfos* (**D**) and *Bdnf-IX* (**E**) gene expression in the CA1 region of the hippocampus 30 minutes after extinction recall. Behaviour: per duration/sex group, n = 8 (total = 48) qPCR: per duration/sex group, n = 8 except 20min and 60min male (n = 7) (total = 46). * = *p* < 0.05; ** = *p* < 0.01; *** = *p* < 0.001.

We showed that in both the 10-minute recall session (48 hours after CFC) and the 2-minute recall session (48 hours after 10min recall), pre-exposure conditions impacted freezing behaviour as an index of CFM, and this differed between males and females (Fig.2B and C). Specifically, during the 10-minute recall session, there was an interaction between Sex and PE duration (*F*_(2,42)_ = 3.501, *p* = 0.032). There was a main effect of PE duration (*F*_(2,42)_ = 9.895, *p* < 0.001), but no main effect of Sex (*F*_(1,42)_ = 0.196, *p* = 0.658). An interaction between Sex and PE duration was also found 48 hours later during the 2-minute recall session (*F*_(2,42)_ = 3.778, *p* = 0.031). In this session there was no main effect of PE duration (*F*_(2,42)_ = 2.056, *p* = 0.141) or Sex (*F*_(1,42)_ = 0.134, *p* = 0.716). In both recall sessions, these interactions manifested as a reduction in conditioned freezing behaviour, and greater LI, with increasing durations of pre-conditioning context pre-exposure in males, but not females. Notably, in females the highest levels of freezing, and least LI, were seen in the 20min PE group, particularly during the first recall session.

Thirty-minutes after the second recall session the expression of the immediate early genes *Cfos*, a marker of neuronal activity, and *Bdnf-IX*, which is known to regulate synaptic plasticity were measured in the CA1 region of the hippocampus using qPCR. *Cfos* and *Bdnf-IX* were therefore selected to determine whether the sex-dependent behavioural effects of PE duration were accompanied by differences in hippocampal neuronal activation and synaptic plasticity, respectively.

There was an interaction between Sex and PE duration in *Bdnf-IX* (*F*_(2,40)_ = 3.271, *p* = 0.048) and *Cfos* expression (*F*_(2,40)_ = 5.796, *p* = 0.006) (Fig.2D and E). In *Bdnf-IX* expression, there was no main effect of Sex (*F*_(1,40)_ = 0.250, *p* = 0.620) or PE Duration (*F*_(2,40)_ = 0.222, *p* = 0.802). In *Cfos* expression, there was a main effect of sex with greater expression in females compared with males (*F*_(1,40)_ = 9.438, *p* = 0.004), but no main effect of PE duration (*F*_(2,40)_ = 2.332, *p* = 0.110). Thus, as was found in the behavioural measures, we showed that for both *Bdnf-IX* and *Cfos* expression, PE duration impacted gene expression in the CA1 region of the hippocampus in a manner that differed between sexes.

Together, these behavioural and molecular findings suggest that the relationship between pre-exposure duration and strength of the LI effect differs between sexes. The existing literature on latent inhibition indicate that longer pre-exposure durations result in reduced expression of associative fear memories[9,150]. The data here show that to be the case in males, but not in females, highlighting that the findings of research conducted primarily in males are not necessarily generalisable to females. The *Cfos* data, which provides an indirect measure of neuronal activity, broadly mirrored the behavioural observations consistent with a role for hippocampal processing and the expression of LI in both males and females. In contrast, *Bdnf-IX* expression showed an inverse pattern to the behavioural response. This divergence suggests that BDNF-related plasticity mechanisms may not directly parallel the behavioural expression of fear but may instead reflect a distinct role for BDNF-related mechanisms in regulating LI and expression[151]. Nevertheless, the interaction between sex and PE duration observed in both *Cfos* and *Bdnf-IX* expression after fear memory recall indicates that the hippocampal molecular response to PE differs between the sexes. These findings are consistent with the possibility that sex-dependent differences in hippocampal processing and synaptic plasticity contribute to the expression of LI. The precise relationship between these molecular changes and the mechanisms underlying LI and fear memory expression remain to be established.

## 6. Discussion

Despite several decades of study of latent inhibition, few studies have attempted to characterise sex differences in LI. Across both human and rodent LI studies, the majority of experiments did not reveal sex differences in LI. However, this was not a universal finding, and in a number of studies, sex differences were shown in the relationship between additional variables and LI. For instance, in humans several studies showed sex differences in the relationship between schizotypy in LI, albeit without a consistent direction of effect. In rodents, this was more extensively assessed, showing sex differences in the effects of factors such as juvenile handling and drug administration on LI. Furthermore, several rodent experiments observed differences in LI in females across phases of the oestrous cycle, or between ovariectomized females with and without oestrogen replacement. Thus, sex effects in LI may be related to a modulatory role of oestrogen on memory processing. No studies found in humans assessed the effects of sex hormones, and a lack of consideration of oestrous may contribute to sex differences not being shown in many studies. Taken together, these findings suggest that although LI may be comparable between sexes at baseline, males and females may differ in the response to particular environmental, pharmacological and hormonal variables. Future research is therefore needed to determine the mechanisms underlying these differential responses. Given the established role of the striatum, hippocampus, mPFC and NAc in LI, investigating whether sex differences in the functioning of the wider neural system contributes to the differential susceptibility to LI disruption will help to improve understanding of the underlying mechanisms. Furthermore, future studies are needed to ascertain the extent to which these differences are attributable to effects of oestrous.

Discrepancies in the observation of sex differences in LI between human and animal studies may reflect differences in experimental paradigms used across species. Owing to important ethical considerations in human LI studies and particularly those including patients, the stimuli used generally bear a neutral valence (e.g., letters on a screen or a flashing light). The use of standardised tasks between studies, also adapted for neuroimaging, allows for replicability and facilitates comparisons of the neural mechanisms underpinning LI. However, psychotic experiences often bear a negative valence and are fearful to the individual[152]. Fear conditioning and the formation of fear associations involve a neuronal circuitry that includes the amygdala, hippocampus and mPFC[153–155]. Thus, there is a possibility that sex differences in fear circuitry relevant to the formation of hallucinations and delusions is not adequately captured by the current array of LI tasks commonly used in humans. Further investigations are required to establish whether sex differences in the fear-related circuitry influence LI and whether these mechanisms are relevant to psychosis. The use of rodent models provides an opportunity to investigate LI using a wider range of conditioning paradigms than can be used in humans, including those that assess the LI of associative fear memory formation and can help determine whether sex differences depend on the valency of learning.

A further factor which likely drives some of the inconsistencies observed between studies are the poor practices in formally testing sex interactions, rather than a true absence of an effect causing many of the null findings. Although many further studies were found which included both male and female animals in their experiments, they did not meet the inclusion criteria as these studies only conducted analyses on sex-aggregated data, without reporting whether sex effects or interactions were tested. Such statistical designs not only limit inferences related to sex differences, but reduce the overall interpretability of results, as false negatives may occur in main effects if variables have opposing directions of effects between sexes, or statistical power will be diluted if they exert effects in only one sex. A greater number of studies testing and reporting these interactions will help to address the still open question of whether sex meaningfully moderates LI in psychosis-spectrum cohorts.

LI may be also affected by sex differences in neural processing. The present study provides evidence of this using a behavioural paradigm of hippocampal-dependent contextual fear memory formation and recall across varying PE durations and both behavioural and molecular outputs. Females showed a different sensitivity to PE duration a with corresponding differences in hippocampal molecular responses following memory recall. In the literature review, only a small minority of the studies investigated LI of contextual fear conditioning, rather than cued fear conditioning. Given the involvement of the hippocampus in contextual associative memory and evidence of sexual dimorphism in this structure, LI assessed using CFC may be particularly sensitive to sex differences. Indeed, the present work showed interactions between sex and PE duration, a finding which is reflected by concurrent interactions between sex and pre-exposure duration in behavioural measures and corresponding interactions the expression of *Cfos* and *Bdnf-IX*, markers associated with neuronal activity and synaptic plasticity in the CA1 region of the hippocampus respectively. These findings suggest that the hippocampal processes through which pre-exposure information influences subsequent associative learning may differ between sexes. Future work is required to test this hypothesis and could extend the approach to a wider range of brain regions (e.g., NAc, mPFC and amygdala), and investigate a wider range of behavioural paradigms, also explicitly considering oestrous phase. Also, because the molecular findings in the present study were obtained following fear memory recall, further assessment is required to establish the neurobiological mechanisms of the encoding of contextual information during pre-exposure, and how this information is used during conditioning.

Aberrant associative learning and memory mechanisms are thought to contribute to the formation of hallucinations and delusions in disorders such as schizophrenia and bipolar disorder. Current pharmacological treatment options for psychosis are plagued with poor efficacy and significant side effects[156]. Hence, it is crucial that novel precision treatment options that specifically target biologically causal mechanisms are developed. Recent transcriptomic analyses of psychosis GWAS data have revealed enrichments in genes highly expressed in the hippocampus as well as those involved in synaptic organisation, differentiation and transmission[157]. Hence, the hippocampus may be a key target for novel pharmacological therapeutics. Developing our understanding of the sexually dimorphic nature of hippocampal-dependent cognition underpinning psychosis and its neurobiological substrates may help elucidate the sex differences in the manifestation and pathophysiology of disorders such as schizophrenia and bipolar disorder. Understanding the specific aetiology of neuropsychiatric disorders in both sexes will then facilitate the development of precision treatment approaches specific to both sexes.

## Acknowledgements

This research was funded by the Medical Research Council, UK, grant no. MR/R011397/1 and legacy funds, with further support from the Hodge Centre for Translational Neuroscience, Cardiff University.

## Data availability

Behavioural and gene expression data included in this article are available at: https://figshare.com/s/6e22bc839e403216c7bf

